# SNR-Efficient Whole-Brain Pseudo-Continuous Arterial Spin Labeling Perfusion Imaging at 7 Tesla

**DOI:** 10.1101/2024.11.12.623276

**Authors:** Joseph G. Woods, Yang Ji, Hongwei Li, Aaron Hess, Thomas W. Okell

## Abstract

**Purpose:** To optimize pseudo-continuous arterial spin labeling (PCASL) parameters to maximize SNR efficiency for RF power constrained whole brain perfusion imaging at 7 T.

**Methods:** We used Bloch simulations of pulsatile laminar flow to optimize the PCASL parameters for maximum SNR efficiency, balancing labeling efficiency and total RF power. The optimization included adjusting the inter-RF pulse spacing (TR_PCASL_), mean B_1_^+^ (B_1 ave_), slice-selective gradient amplitude (G_max_), and mean gradient amplitude (G_ave_). In vivo data were acquired from four volunteers at 7 T to validate the optimized parameters. Dynamic B_0_-shimming and flip angle adjustments were utilized to avoid needing to make the PCASL parameters robust to B_0_/B_1_^+^ variations.

**Results:** The optimized PCASL parameters achieved a significant (3.1x) reduction in RF power while maintaining high labeling efficiency. This allowed for longer label durations and minimized deadtime, and resulted in a 126% improvement in SNR efficiency in vivo compared to a previously proposed protocol. Additionally, the static tissue response was improved, reducing the required distance between labeling plane and imaging volume.

**Conclusion:** These optimized PCASL parameters provide a robust and efficient approach for whole brain perfusion imaging at 7 T, with significant improvements in SNR efficiency and reduced SAR burden.

## 1. Introduction

Arterial spin labeling^1,2^ (ASL) is a completely non-invasive MRI method that can be used for imaging and quantifying tissue perfusion without the use of injected contrast agents or ionizing radiation. However, the SNR of ASL measurements is inherently limited by blood flow rates and T_1_ relaxation of the labeled blood water.

To achieve sufficient SNR for accurate quantification of brain gray matter (GM) perfusion at 3 T, a relatively low spatial resolution is used (typically ∼3.5x3.5x5 mm^3^) and many images are acquired for signal averaging (typically 25-30^3,4^). In order to attempt to quantify white matter (WM) perfusion**\***, or to use higher spatial resolution readouts, the number of averages must be appropriately increased to maintain sufficient SNR.^5^ For example, scan time would need to be increased by a factor of 16 to balance a factor of 4 reduction in signal. Alternatively, ultra-high field (UHF) MRI provides greater polarization and longer T_1_ relaxation rates, with a potential 2-4 times higher ASL perfusion SNR at 7 T than 3 T.^6^ Due to the increasing availability of 7 T scanners, this represents an attractive approach for probing WM perfusion or using higher spatial resolutions without onerous scan times.

However, ASL faces a number of challenges at UHF, including increased specific absorption rate (SAR) and more inhomogeneous B_1_^+^ and B_0_ fields. Pseudo-continuous ASL (PCASL) is an attractive labeling approach for 7 T because of its ability to generate long label durations with the commonly used, spatially limited, head-only transmit coils.^6^ Unfortunately, PCASL is particularly sensitive to off-resonance and has a high SAR burden, making its translation to 7 T challenging. While previous studies have introduced methods for mitigating the effects of inhomogeneous B_1_^+^ and B_0_ fields on labeling efficiency, the high SAR burden of PCASL has garnered less attention. This SAR burden is, nevertheless, practically limiting because it requires deadtime to be inserted into each TR to stay within SAR limits; this deadtime reduces SNR efficiency and, thus, limits the advantage of acquiring data at 7 T.

Several previous works^10,11,19^ have used variable-rate selective excitation^20,21^ (VERSE) to reduce the power of the PCASL RF pulses. Since these pulses have short durations (∼500 µs), there is very little distortion of the slice profile due to off-resonance phase accrual, even with the large off-resonances encountered at 7 T.^11^ Boland et al.^19^ additionally increased the RF duty cycle to further reduce RF power, by maximizing the duration of the PCASL RF pulse within the inter-pulse spacing (referred to as TR_PCASL_ here) subject to gradient hardware limits. However, these studies used PCASL labeling settings designed for 3 T, which are expected to be less than optimal at 7 T. Although several studies^12,22^ have generated 7 T specific PCASL labeling settings, the settings were chosen to maximize labeling efficiency across a large range of off-resonance and B_1_^+^, ignoring the effect of the SAR burden on the achievable TR and SNR efficiency.

In this work, we use Bloch simulations of pulsatile laminar flow to optimize the PCASL pulse train parameters (Figure 1) to instead maximize SNR efficiency, where the minimum TR is assumed to be SAR constrained (i.e. the minimum TR is proportional to the RF energy deposited within each TR). Instead of aiming to make the PCASL labeling process inherently insensitive to B_0_ and B_1_^+^ variation, as in previous works, we directly correct for these effects at the labeling plane, releasing degrees of freedom which can instead be used to maximize SNR efficiency. Finally, we also explore whether reducing the RF power of the adiabatic inversion pulses used for background suppression (BGS) can further increase SNR efficiency.

**Figure 1:**
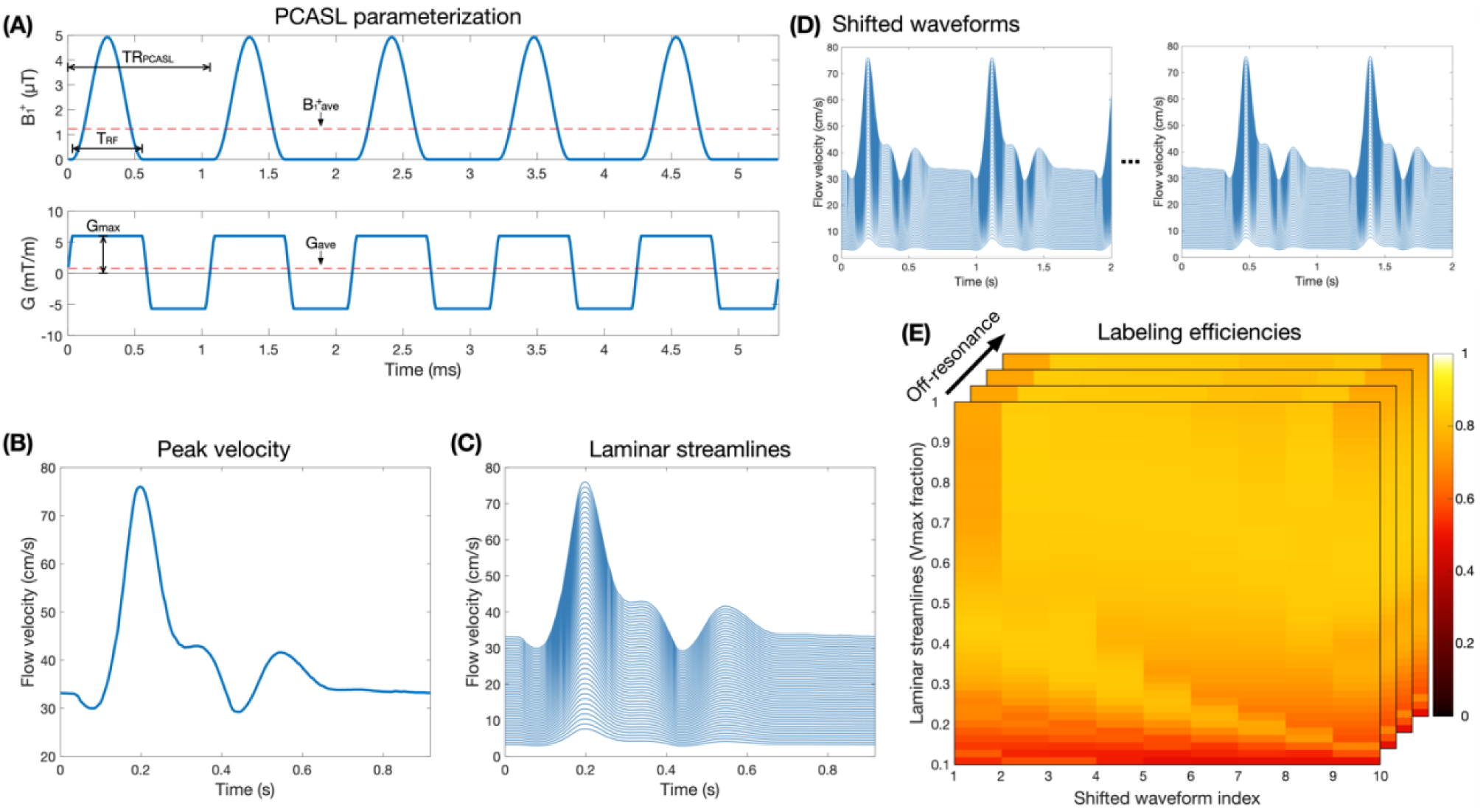
(A) The parameterization of the PCASL RF (top) and gradient (bottom) pulse train. (B) The pulsatile flow velocity waveform, with peak of 76cm/s and minimum velocity of 30cm/s, used for labeling efficiency Bloch simulations, with the simulated laminar streamlines in (C) and the shifted velocity waveforms in (D). (E) shows example results of the labeling efficiency Bloch simulations across laminar streamlines, shifted waveforms, and off-resonance values.

## 2. Methods

### 2.1. PCASL parameter optimization

The PCASL pulse train can be parameterized by 5 variables: the RF pulse duration (T_RF_), the inter-RF pulse spacing (TR_PCASL_), the mean B_1_^+^ (B_1_^+^_ave_), the gradient amplitude during the RF pulse (G_max_), and the mean gradient amplitude (G_ave_) (Figure 1A).^9^ To optimize these parameters to maximize SNR efficiency, we used numerical Bloch simulations of flow-weighted pulsatile laminar flow to simulate the labeling efficiency as described below.

We implemented a custom fixed-duration minimum-SAR VERSE^20,21^ algorithm and applied it to the PCASL pulses in the optimization. Briefly, the PCASL B_1_^+^ and gradient waveforms are transformed to generate a constant amplitude B_1_^+^ pulse with fixed duration, subject to maximum gradient limits. Gradient amplitude and slew rate violations are then repeatedly resolved,^20^ followed by a pulse duration adjustment, until no violations remain.

The evaluated PCASL parameter values were: TR_PCASL_=[0.5, 0.6, 0.7, 0.8, 1.06] ms (TR_PCASL_ between 0.8-1.06 ms are not possible with our scanner due to mechanical resonances), B_1_^+^ _ave_ = 0.1-2.0 µT at 0.1 µT increments (equivalent flip angles of 1.6°-32° for TR_PCASL_ = 1.06 ms), G_max_ = 3.0-15.0 mT/m at 0.5 mT/m increments, and Gave = 0.0-2.0 mT/m at 0.1 mT/m increments. T_RF_ was fixed at 0.5 · TR_PCASL_, i.e., 50% RF duty cycle. Parameter combinations that exceeded gradient hardware limits (max amplitude 80 mT/m, max slew rate 200 T/m/s) were excluded.

Spin isochromats (referred to simply as “spins”) were simulated by numerically integrating the Bloch equations with the hard pulse approximation using a time-step of 10 µs, with T1 = 2.1 s^23^ and T2 = 0.06 s^24^ assumed for blood at 7 T. To simulate the labeling efficiency, spins were simulated moving in one dimension from 5 cm below to 8 cm above the labeling plane, similar to Zhao et al.^17^ The final longitudinal magnetization, Mz, was corrected for T_1_ relaxation with the labeling efficiency then calculated as 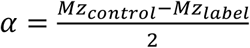.

The pulsatile velocity waveform from Zhao et al.^17^ was used to describe the peak velocity in a laminar flow profile, with minimum and maximum velocities of 30 cm/s and 76 cm/s (Figure 1B). However, unlike Zhao et al.,^17^ who simulated plug flow for a range of velocities and then took the flow-weighted mean of the calculated labeling efficiencies across the velocity distribution, we directly simulated the spins moving according to the pulsatile laminar flow waveform.

Fifty laminar streamlines were simulated, equally spaced from 10% to 100% of the maximum velocity waveform (Figure 1C) (streamlines less than 10% of the maximum contribute little to the flow-weighted average, but the simulations take an increasingly long time). Because the labeling efficiency is affected by the particular velocity at which spins are moving when they cross the center of the labeling plane, we also shifted the velocity waveform by 10 equal increments and simulated the 50 laminar streamlines for each case (Figure 1D). Finally, to ensure robust labeling efficiency over a small range of off-resonance (discussed in section 2.2), 11 values, evenly spaced between −50 Hz and 50 Hz, were simulated for each case (Figure 1E). The final labeling efficiency for each set of PCASL parameters was then given by the flow-weighted mean:

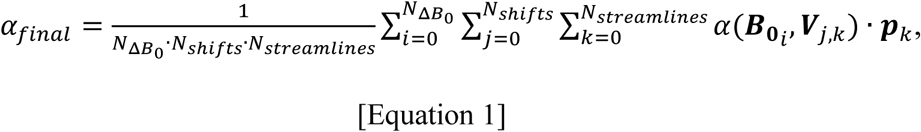

where **V** is the velocity waveform array and 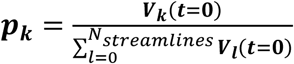 is the normalized flow-weighted laminar probability distribution function.

Aliased labeling planes can perturb the static tissue signal in the imaging volume if the condition 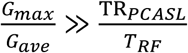 is not met.^9^ To more informatively guide the choice of G_max_ and G_ave_, Zhao et al.^22^ proposed placing the first aliased labeling plane at the 3rd zero-crossing of the Fourier transform of the PCASL RF envelope by setting 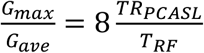. Here, we instead simulate the effects of aliased labeling planes on static tissue for each parameter combination. These Bloch simulations included stationary spins located from −16 to 16 cm from the labeling plane, spaced every 0.2 mm, under the influence of a 1800 ms labeling train, using T1 = 2.1 s, T2 = 0.06 s, and a time-step of 10 µs. The control-label Mz difference was then convolved with the simulated slice profile of a 90° Hann-windowed sinc excitation pulse (duration=2.56 ms, time-bandwidth product=3.2, slice thickness=5 mm) to simulate the effect that the aliased labeling planes have on the acquired 2D images (see section 2.3). PCASL parameter combinations that had a static tissue perturbation of more than 0.1% of |*M*_0_| beyond 1.8 cm from the labeling plane were excluded from the optimization. Note, 1.8 cm was the mean distance between the labeling plane and the bottom of the cerebellum in 6 previously scanned healthy volunteers.

The optimally SNR efficient set of PCASL parameters, θ_opt_, were then found by maximizing the simulated SNR efficiency, i.e.,:

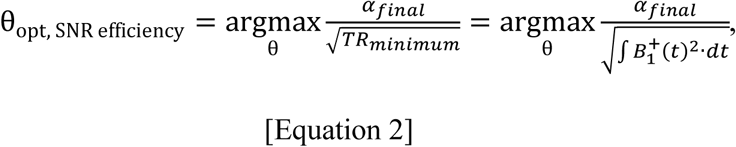

since the minimum TR is typically proportional to the total RF energy deposited during each TR in 7 T PCASL sequences,^25^ i.e., 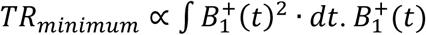is the RF amplitude waveform of a whole TR.

For comparison, we also chose the set of parameters that maximized SNR, ignoring RF power, which is equivalent to maximizing the labeling efficiency:

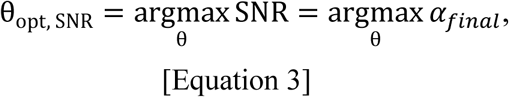

### 2.2. Off-resonance correction at the labeling plane

Although unbalanced PCASL^9^ (G_ave_ = 0 in the control condition) is intrinsically more robust to off-resonance effects,^17^ it has mis-matched eddy currents between label and control conditions that may result in artifacts,^26^ and it is not compatible with vessel-encoded labeling,^27^ which we plan to use in future applications. Therefore, we used balanced PCASL,^27^ where the only difference between label and control conditions is that the PCASL RF pulses in the control condition have an additional π phase added to every other pulse.

To minimize off-resonance at the labeling plane, and to avoid needing to make the PCASL pulse train itself insensitive to B_0_ variation at the expense of increased SAR, we used dynamic linear B_0_-shimming of the labeling plane, as described by Ji et al.^15^ That is, after performing 2nd order B_0_-shimming of the imaging volume, the linear shims were dynamically adjusted in real-time during the sequence so that the 4 feeding arteries at labeling plane were optimally shimmed during PCASL labeling, with the shim values being reset to the imaging volume optimal values at the end of the labeling train.

Ji et al.^15^ demonstrated that off-resonance at the labeling plane could be robustly reduced to less than ±50 Hz, greatly improving labeling efficiency and only requiring the PCASL parameters to be robust to ±50 Hz variation in the main magnetic field. Dynamic B_0_-shimming was used for all protocols in this work, including the literature PCASL settings described below.

### 2.3. In vivo data acquisition

Four volunteers (one female, age 22-36 years) were recruited and scanned under a technical-development protocol with approval from local ethics and institutional committees. Data were acquired with a MAGNETOM 7T Plus scanner (Siemens Healthineers, Forchheim, Germany) equipped with an 8Tx/32Rx head coil (Nova Medical, Wilmington, MA, USA). All data were acquired in circularly polarized mode. The volunteers were asked to lie still during the scan but were not required to stay awake.

#### 2.3.1. Auxiliary scans

A multi-slab 3D time-of-flight scan (0.26x0.26x0.6 mm^3^) was used to place the transverse PCASL labeling plane at the middle of the V3 section of the vertebral arteries and locate the center of the four feeding arteries.

A 3DREAM^28^ B_1_^+^-mapping sequence (5x5x5 mm, duration=6 s), covering the brain and neck, was performed to calibrate the transmit voltages. The PCASL sequence reference voltage, *V*_*reference*_, was set to the mean value within the brain at the central slice. The nominal PCASL RF flip angle was then increased to account for the lower B_1_^+^ at the labeling plane by a factor 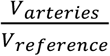, where *V*_*arteries*_ is the mean reference voltage of the four feeding arteries at the labeling plane.

A single slice 2D dual-echo GRE fieldmap (0.73x0.73x3 mm^3^, ΔTE=1.02 ms, flip angle=30°, TR=30 ms, duration=18 s) was acquired with an identical B_0_ shim as the PCASL data for calculating the optimal in-plane linear shim adjustments and inter-pulse phase correction for the PCASL labeling, as described in Ji et al.^15^. An MPRAGE^29^ scan (0.7x0.7x0.8 mm_3_, TI=1100 ms, TR=2.6 s, flip angle=5°) was acquired for image registration purposes and tissue segmentation. A 24-slice 2D dual-echo GRE fieldmap (1.72x1.72x5 mm^3^, ΔTE=1.02 ms), with matched coverage and identical B_0_ shim to the PCASL data, was used for distortion correction of the PCASL EPI data.

#### 2.3.2. PCASL perfusion acquisition

In addition to acquiring perfusion data with the optimized maximum SNR efficiency and maximum labeling efficiency PCASL parameters, we also included the PCASL parameters from Zhao et al.,^22^ a recent 7 T PCASL study that used PCASL parameters optimized for robustness against large variations in B_1_^+^and B_0_ (Table 1). Although Zhao et al. did not use VERSE, we acquired data with and without VERSE applied to the PCASL RF pulses to demonstrate the SNR efficiency advantages.

**Table 1:** PCASL parameters, relative PCASL RF power, and simulated labeling efficiency and SNR efficiency. The “nominal” RF flip angle and B_1_^+^_ave_ is relative to sequence reference voltage, which is typically calibrated within the imaging volume, whereas the “in vivo B_1_^+^” values are from simulations where the mean B_1_^+^ at the labeling plane across the four feeding arteries in 4 volunteers was used. The PCASL RF voltage was adjusted for the max SNR efficiency and max labeling efficiency protocols in vivo to achieve the intended B_1_^+^_ave_ at the labeling plane, so the labeling efficiency for these protocols remains unchanged with the in vivo B_1_^+^ values, whereas the PCASL RF voltage was not adjusted for the literature protocols, in keeping with the setup in ref ^22^.

| PCASL Parameters | RF flip angle<br>nominal $B_1^+$ /<br>in vivo $B_1^+$ | $B_{1^+ave}$<br>nominal $B_1^+$ /<br>in vivo $B_1^+$ | $T_{RF}$ | $TR_{PCASL}$ | $G_{max}$ | $G_{ave}$ | Labeling<br>efficiency<br>nominal<br>$B_1^+$ / in<br>vivo $B_1^+$ | Relative<br>PCASL<br>RF power<br>nominal<br>$B_1^+$ / in<br>vivo $B_1^+$ | Relative<br>sequence<br>SNR<br>efficiency<br>nominal $B_1^+$<br>/ in vivo $B_1^+$ |
| --- | --- | --- | --- | --- | --- | --- | --- | --- | --- |
| Max SNR<br>efficiency | 9.7° | 0.6 μT | 530<br>ms | 1060 ms | 5.5<br>mT/m | 0.2<br>mT/m | 0.72 / 0.72 | 1.0 / 1.0 | 1.78 / 1.55 |
| Max<br>labeling<br>efficiency | 15.9° | 1.3 μT | 400<br>ms | 800 ms | 8<br>mT/m | 0.5<br>mT/m | 0.82 / 0.82 | 4.9 / 4.9 | 1.42 / 1.00 |
| Literature | 15° / 8.2° | 1.72 μT / 0.94<br>μT | 300<br>ms | 570 ms | 5.9<br>mT/m | 0.4<br>mT/m | 0.76 / 0.79 | 10.4 / 3.1 | 1.00 / 1.16 |
| Literature<br>with<br>VERSE | 15° / 8.2° | 1.72 μT / 0.94<br>μT | 300<br>ms | 570 ms | 5.9<br>mT/m | 0.4<br>mT/m | 0.76 / 0.79 | 8.1 / 2.4 | 1.10 / 1.28 |

Due to differences in real-time SAR monitoring, we were not able to use Zhao et al.’s^22^ exact parameters; specifically, we increased TR_PCASL_ from 550 µs to 570 µs. Additionally, we rounded G_max_ from 5.867… mT/m to 5.9 mT/m. These changes resulted in negligible differences in simulated labeling efficiency, as shown in **Figure S1**.

PCASL perfusion data were acquired with the four sets of PCASL parameters listed in Table 1. The sequence consisted of a slab-selective (thickness 160 mm) water suppression–enhanced– through–T_1_^+^-effects^30,31^ pre-saturation module covering the entire brain, followed immediately by the PCASL labeling module and a 1800 ms post-label delay (PLD). The vendor implemented EPI readout was used for data acquisition: 24 slices, 5 mm slice thickness, 1 mm slice separation, FOV=220x220 mm^2^, voxel size=3.44x3.44 mm^2^, flip angle=90°, 6/8 phase partial Fourier, TE=13 ms, and bandwidth=2004 Hz/pixel. The 2nd order B_0_ shim was optimized for the imaging region only.

During the PLD, two non-spatially-selective hyperbolic secant (sech) adiabatic inversion pulses were used to null spins with T_1_ = 1000 ms and 2000 ms, 100 ms before the first readout using the formula in ref ^32^. Given a pulse duration of 10.24 ms, amplitude truncation of 4% (β = 763.99 rad/s),^33^ and B_1_^+^_max_ = 20 µT, the shaping factor, µ, was optimized to maximize the inversion efficiency over a B_1_^+^ variation of ±50% and B_0_ variation of ±500 Hz, sampled every 1% and 10 Hz, respectively. This resulted in µ = 7.06.

The longest PCASL label duration, up to a maximum of 1800 ms, was set subject to the coil manufacturer’s 1 second RF power limit (max 70 W average). The TR was then set to the minimum value possible based on the scanner predicted 1st level SAR value and the number of averages was set to achieve a scan duration of 4 minutes per protocol, not including the M_0_ calibration image.

An additional high spatial resolution data set (1.95x1.95x4 mm^2^) was acquired in one volunteer using the optimized max SNR efficiency PCASL parameters to demonstrate data quality achievable using 7T PCASL with these settings. The label duration and PLD were both 1800 ms. The EPI readout settings were: 29 slices, 4 mm slice thickness, 0.8 mm slice separation, FOV=250x250 mm^2^, flip angle=90°, 6/8 phase partial Fourier, in-plane GRAPPA acceleration factor=2, TE=13 ms, bandwidth=2055 Hz/pixel, and scan time=4 minutes. A FOV matched 29-slice 2D dual-echo GRE fieldmap (1.95x1.95x4 mm^3^, ΔTE=1.02 ms) was also acquired for EPI distortion correction.

### 2.4. Reduced SAR background suppression

In addition to the SNR efficiency optimization of the PCASL settings, we also investigated whether reducing the SAR of the sech BGS inversion pulses could improve the SNR efficiency of the sequence.

We applied our minimum SAR VERSE algorithm to the sech pulse, optimizing µ for the reformatted RF pulse as described above. SAR reduction was restricted to 25% to limit the reduced off-resonance performance seen at higher SAR reduction levels.^34^ The phase waveform of the VERSE sech pulse was then further optimized to maximize inversion efficiency.

In vivo data in the same volunteers and with similar acquisition parameters to the standard resolution data was acquired to evaluate whether the SNR efficiency of the sequence was improved by this approach. Full methods details are provided in Supporting Information 1.

### 2.5. Postprocessing

The MPRAGE images were processed using FSL’s (version 6.0.6.5) fsl_anat script, which performs brain extraction^35^, B_1_^+^ bias-field correction, and tissue segmentation.^36^

The PCASL data was processed and quantified using oxasl^37,38^ (https://github.com/physimals/oxasl, version 0.2.1.post17), which performs affine motion correction,^39–41^ pairwise subtraction of label and control data, EPI distortion correction, and registration of the structural and PCASL data. Perfusion quantification used the model in Alsop et al.^3^ with T_1_^+^ = 2.1 s and BGS inversion pulse efficiency = 0.944 and 0.931 for the sech and the phase-optimized VERSE sech pulse, respectively (derived from Bloch simulations for ΔB_1_^+^ = ±50%, ΔB_0_ = ±500 Hz, T_1_^+^ = 2.1 s, T_2_ = 0.06 s). Gray matter masks were created by thresholding the partial volume maps at 0.5.

SNR efficiency maps were generated as 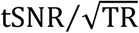. The gray matter masks were then used to calculate the mean SNR efficiency for each subject.

## 3. Results

### 3.1. PCASL parameter optimization

Of the 52500 simulated parameter combinations, 23040 (43.9%) were not simulated because they exceeded gradient slew rate limits. A further 26839 (51.1%) parameter combinations were excluded because they had a static tissue perturbation of more than 0.1% of |*M*_0_| beyond 1.8 cm from the labeling plane. This left 2621 (5%) of the parameter combinations that were included in the labeling efficiency and SNR efficiency optimization. Histograms of the included PCASL parameters are shown in Figure S2, demonstrating that a reasonable range of the parameters were still included. Additionally, Figure S3(A-C), demonstrates the optimal max SNR efficiency parameters had a theoretical SNR efficiency only 3% lower than the unconstrained maximum.

The resulting optimal max SNR efficiency and max labeling efficiency PCASL parameters are provided in Table 1, with the B_1_^+^ and gradient waveforms shown in Figure 2. Table 1 also reports the simulated mean labeling efficiencies across a range of ±50 Hz off-resonance, the relative PCASL RF power, and the mean relative SNR efficiency (accounting for the total sequence RF power) for each protocol. Each of these values is reported for the nominal PCASL B_1_^+^_ave_, and for the B_1_^+^_ave_ achieved in vivo at the labeling plane for the four volunteers, where the nominal PCASL flip angle was increased to achieve the intended B_1_^+^_ave_ for the max SNR efficiency and max labeling efficiency protocols.

**Figure 2:**
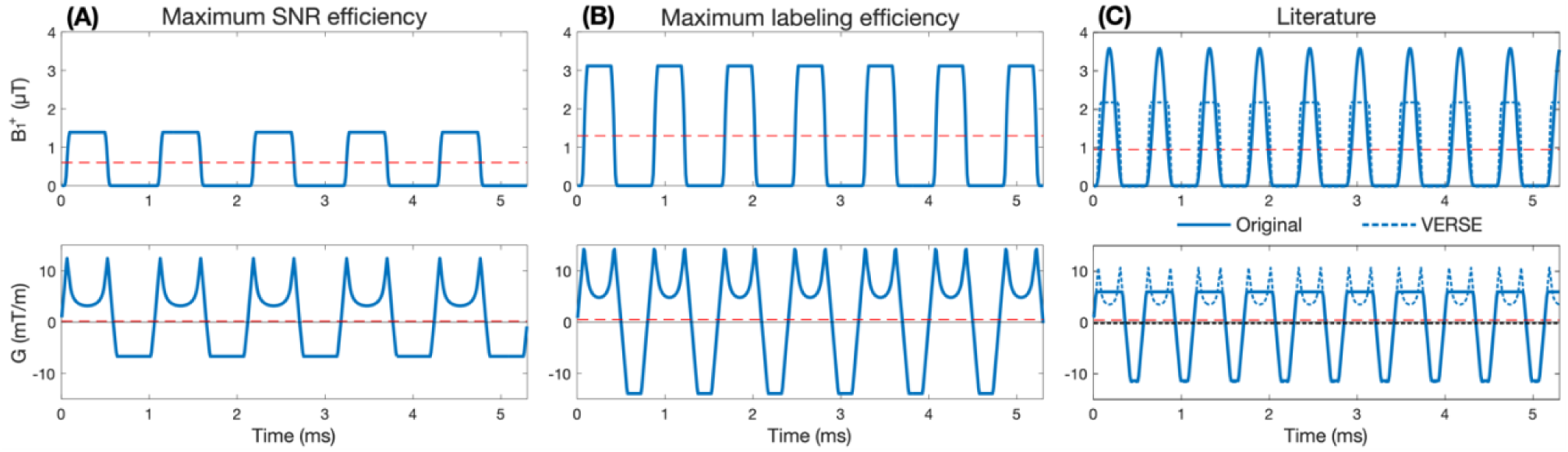
The PCASL B_1_^+^ and gradient waveforms (blue lines) for the PCASL parameters listed in Table 1. B_1_^+^_ave_ and G_ave_ are shown as red dashed lines. The literature PCASL waveforms with VERSE are shown in (C) as blue dashed lines. The literature waveforms shown in (C) used the mean B_1_^+^_ave_ achieved in vivo at the labeling plane across the four subjects.

From Table 1 we can see that the max SNR efficiency parameters used a low B_1_^+^_ave_, long TR_PCASL_, low G_max_, and low G_ave_ to achieve 2.4-4.9 times lower in vivo PCASL SAR, while only having 8.9%-12.2% lower labeling efficiency. This meant that the theoretical SNR efficiency of the max SNR efficiency protocol is 22%-55% higher in vivo than the three comparison protocols.

To further understand the relative SNR efficiency of each protocol, Figure 3(A-B) shows the simulated SNR efficiencies and labeling efficiencies for a range of constant velocities (mean across ±50 Hz off-resonance range). These results reflect those in Table 1, but also demonstrate that the max SNR efficiency protocol has higher SNR efficiency at all velocities between 5-80 cm/s, rather than just those represented in the pulsatile flow waveform.

**Figure 3:**
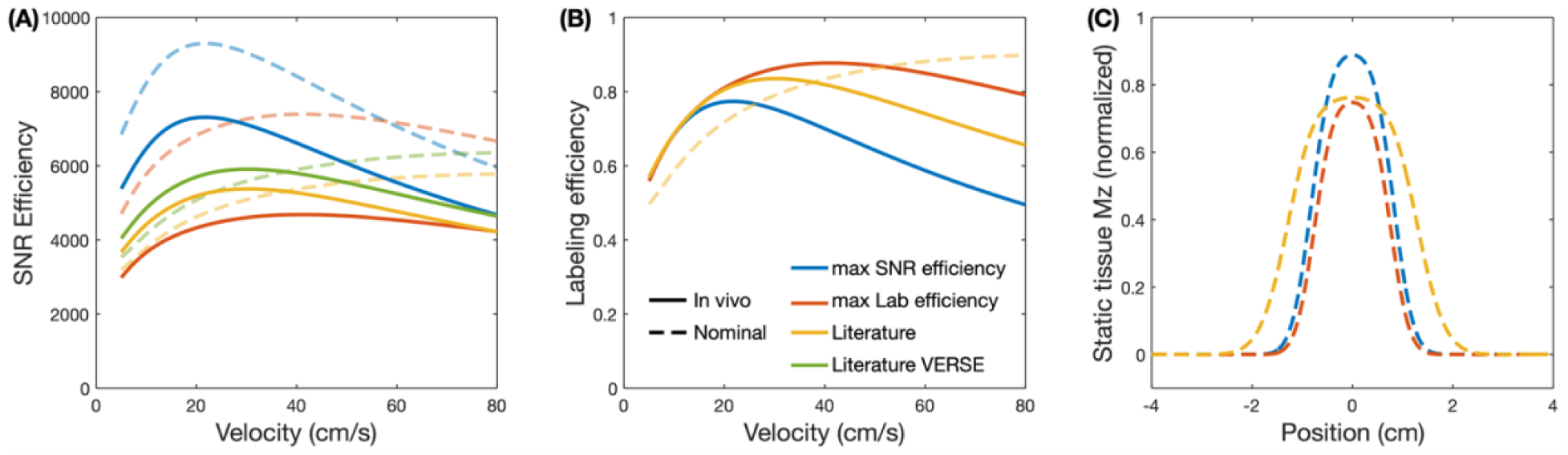
The mean simulated SNR efficiencies (A) and labeling efficiencies (B) across the ±50 Hz off-resonance range for each protocol for a range of constant velocities. In both cases, the data was simulated for each protocol using the nominal and mean in vivo B_1_^+^_ave_, where the SAR of the PCASL pulses relative to the rest of the sequence RF pulses was recalculated to account for the in vivo B_1_^+^ adjustments. The static tissue response for each protocol is shown in (C) assuming the nominal B_1_^+^_ave_.

Despite the low G_max_ and G_ave_ of the max SNR efficiency protocol, the G_max_⁄G_ave_ ratio was very high at 27.5, resulting in a negligible aliased labeling plane response despite the first aliased labeling plane not occurring at a zero-crossing of the PCASL RF Fourier transform. In contrast, the max labeling efficiency parameters used a higher B_1_^+^_ave_, G_max_, and G_ave_, and a shorter TR_PCASL_. The G_max_⁄G_ave_ ratio in this case was 16, meaning the first aliased labeling plane coincided with the 3rd zero-crossing of the PCASL RF Fourier transform, since TR_PCASL_⁄T_RF_ = 2.^22^

In terms of the central lobe of the static tissue response, at ΔMz = 0.001 it extended roughly 1.78 cm, 1.62 cm, and 2.64 cm beyond the center of the labeling plane for the max SNR efficiency, max labeling efficiency, and literature protocols, respectively (Figure 3(C)). It should be noted that for the max SNR efficiency parameters, although the SNR efficiency did not vary much with TR_PCASL_ (<0.3% between 0.5 - 0.8 ms, but 2.2-2.5% lower for TR_PCASL_ = 1.06 ms), principally because the simulated off-resonance range was small, the static tissue central lobe width decreased with increasing TR_PCASL_ (Figure S3(D)), with only TR_PCASL_ = 1.06 ms satisfying the static tissue constraint for the max SNR efficiency parameters.

The use of VERSE with the literature protocol reduced the PCASL RF power by 22% with negligible difference to the simulated labeling efficiency (−0.02%), the latter result being in agreement with previous works.^11,19^ This increased the SNR efficiency of the protocol by 9.9%. For the max SNR efficiency and max labeling efficiency protocols, VERSE reduced the PCASL RF power by 25% and 22%, respectively.

### 3.2. In vivo SNR efficiency comparison

Across the four subjects, the nominal PCASL flip angle for the max SNR efficiency and max labeling efficiency protocols was increased by a factor of (mean ± SD) 1.82 ± 0.07 to achieve the intended B_1_^+^_ave_. Likewise, because this was not adjusted for the literature protocol (as per ref), the effective mean PCASL B_1_^+^_ave_ was 1.82 times lower than the nominal value (flip angle = 8.2° rather than 15°).

The max SNR efficiency PCASL parameters operated under the RF coil manufacturer’s 1 s average transmit power limit for all subjects, meaning a label duration of 1800 ms could be used. However, the max labeling efficiency, literature, and literature with VERSE, protocols exceeded this limit in all subjects, requiring us to greatly reduce the label duration below 1 s. Specifically, the label durations were 275 ± 54 ms, 425 ± 50 ms, and 556 ± 63 ms, respectively (Figure 4(A)), highlighting the high SAR burden of these protocols. The deadtimes added to each TR which then satisfied the 1st level SAR limits are shown in Figure 4(B). Of note, the minimum TR was achieved with the max SNR efficiency protocol (4.7 s) in one subject, but not for the other protocols or subjects.

**Figure 4:**
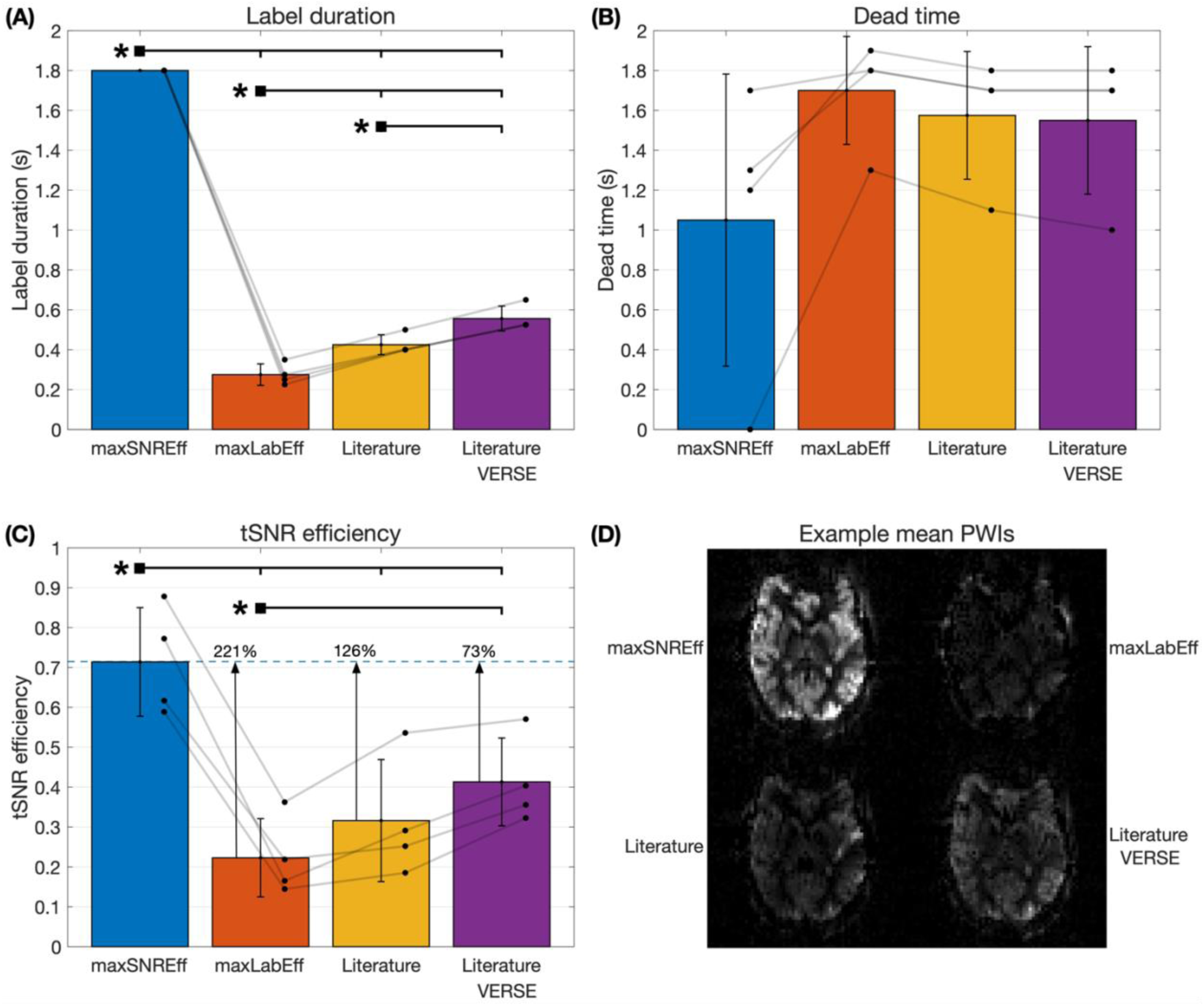
The results of the main in vivo SNR efficiency comparison between the max SNR efficiency (maxSNREff), max labeling efficiency (maxLabEff), literature, and literature with VERSE, protocols. (A) The maximum label durations, up to 1800 ms, achieved for each protocol subject to the coil manufacturer’s 1 s average RF power limit. (B) The minimum added sequence deadtime achieved for each protocol, subject to the 1st level SAR limits. (C) The subject-level mean gray matter SNR efficiencies for each protocol, calculated as the temporal SNR divided by the square-root of the minimum TR. (D) Example minimally processed mean perfusion weighted images for each protocol from one subject, with consistent windowing. In each graph, the bar graphs show the mean and standard deviation across subjects. Individual subject values are plotted as dots. Significant differences, calculated using a paired-sample t-test with Bonferroni correction for 7 comparisons (including the background suppression comparison below), are shown with an asterisk (*).

Figure 4(C) shows the quantitative in vivo gray matter SNR efficiency results, demonstrating that across the four subjects the max SNR efficiency protocol achieved 221%, 126%, and 73% higher SNR efficiency on average than the max labeling efficiency, literature, and literature with VERSE protocols, respectively. The SNR efficiency maps for an example subject are shown in Figure 5 and clearly demonstrates that a much higher SNR efficiency is achieved across the entire brain for the max SNR efficiency protocol. Quantified perfusion maps are shown for the same subject in Figure 6, demonstrating much noisier data for the comparator protocols than the max SNR efficiency protocol.

**Figure 5:**
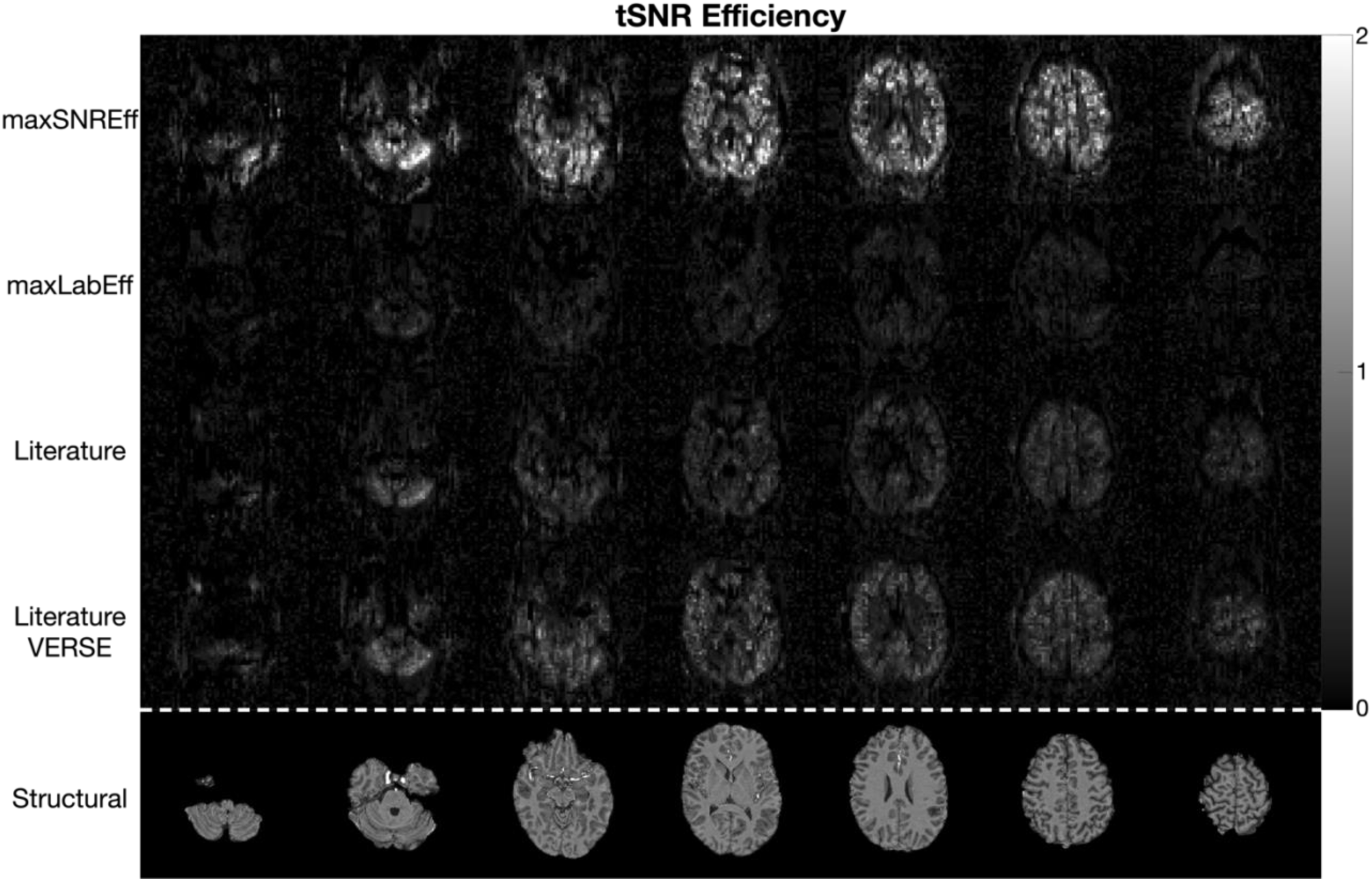
The results of the main in vivo tSNR efficiency comparison in one subject. Each row shows seven slices from each protocol, with the corresponding slices from the MPRAGE structural scan shown in the bottom row.

**Figure 6:**
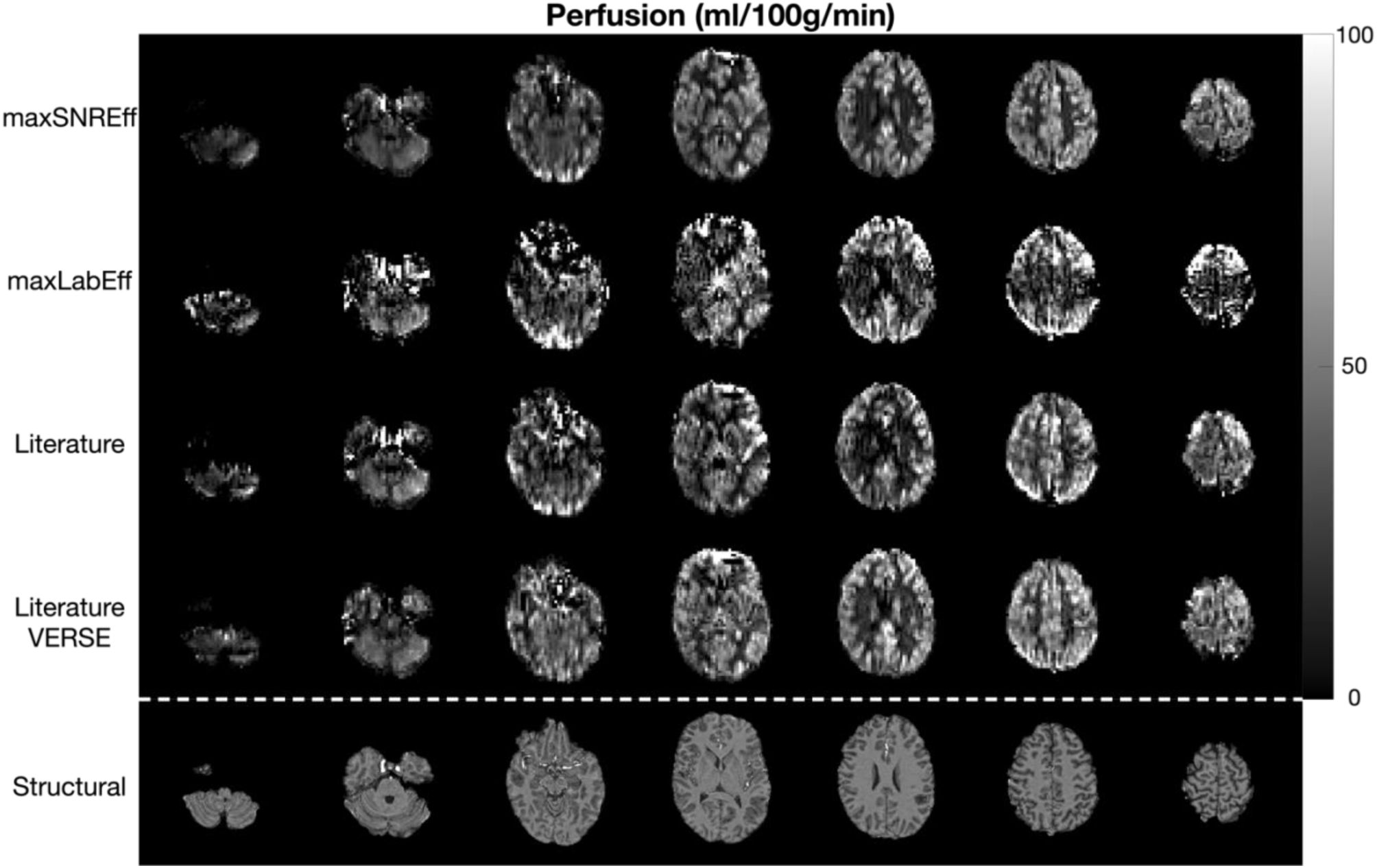
The quantified CBF maps from the main comparison in the same subject as in Figure 5. Each row shows seven slices from each protocol, with the corresponding slices from the MPRAGE structural scan shown in the bottom row.

### 3.3. Reduced SAR background suppression

The B_1_^+^ waveforms and simulated inversion performance for the standard sech pulse, VERSE transformed sech pulse, and VERSE transformed sech pulse with optimized phase waveform are shown in Figure 7. All three pulses achieved good inversion efficiency (mean inversion efficiency = 0.944, 0.927, 0.931, respectively) across the target range (ΔB_1_^+^ = ±50 %, ΔB_0_ = ± 500 Hz). Optimizing the phase waveform of the VERSE sech pulse reduced the oscillations at low B_1_^+^ (Figure S4), and modestly improved the mean inversion efficiency. In simulations, use of the optimized-phase VERSE pulse improved the predicted SNR efficiency of the max SNR efficiency protocol by a modest 2.34%, though this did not account for the reduced BGS performance of the pulse, only the reduced ASL signal due to the pulse’s lower inversion efficiency.

**Figure 7:**
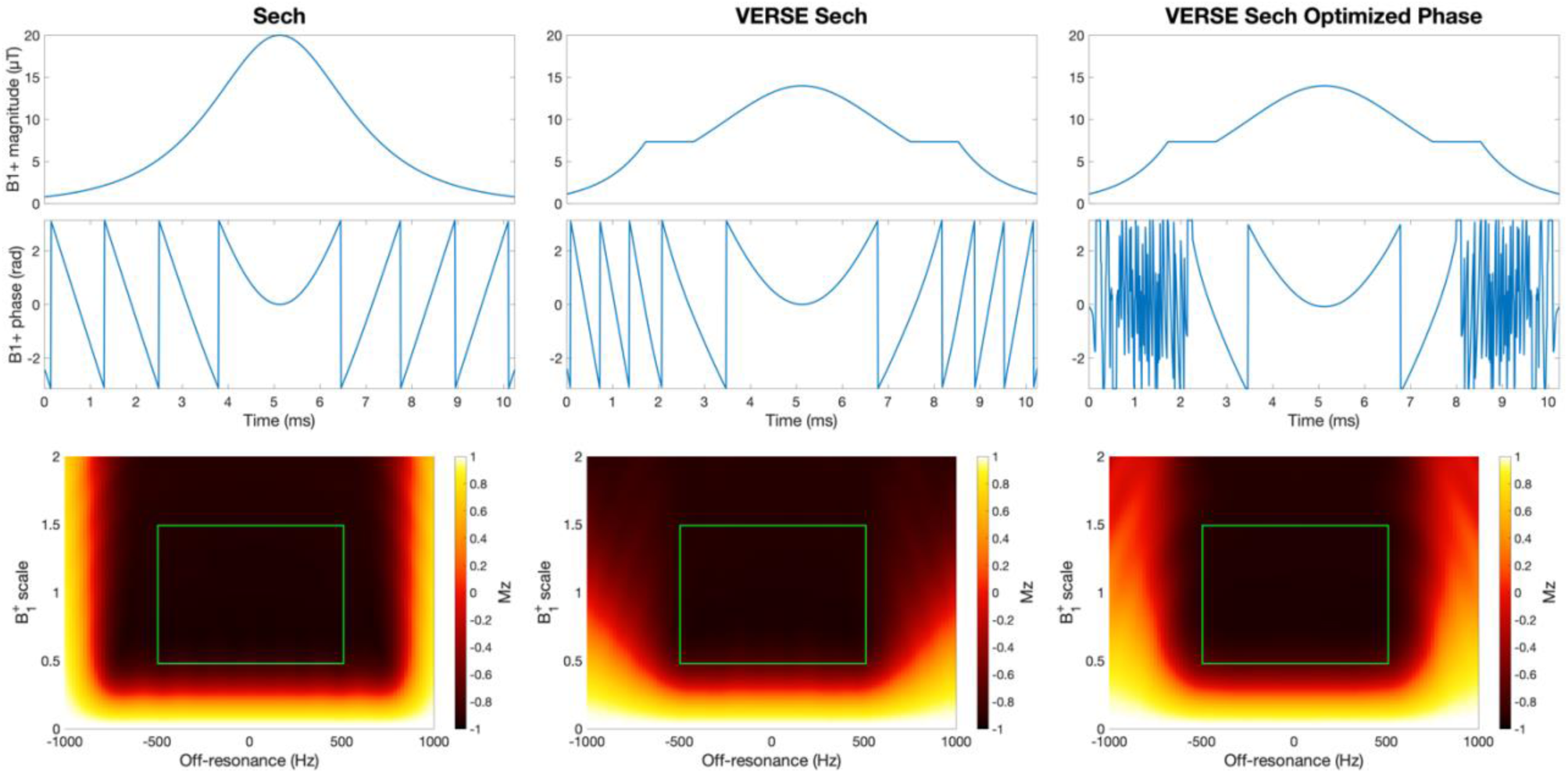
The B_1_^+^ magnitude (top row) and phase (middle row) waveforms and the resulting Mz immediately after the pulse for a wide range of ΔB_1_^+^ and ΔΒ_0_ (bottom row). The green box represents the target ΔB_1_^+^ and ΔΒ_0_ used in the pulse optimizations. Smaller windowing of the Mz plots is shown in Figure S5.

The in vivo inversion efficiency is demonstrated in Figure S5, confirming the marginally lower mean inversion efficiency across the brain for the phase-optimized VERSE sech pulse compared to the standard sech pulse.

The results of the in vivo SNR efficiency comparison between the max SNR efficiency protocol with the standard sech BGS pulse and the optimized-phase VERSE sech pulse are shown in Figure 8 and Figure S6. Although the SAR reduction from the optimized-phase VERSE sech pulse enabled a reduction in TRs of 225 ms, the in vivo tSNR efficiency was 11% lower than when the standard sech pulse was used (0.635 versus 0.714). Nevertheless, the mean GM CBF estimates were similar (42.3 mL/100g/min versus 43.7 mL/100g/min).

**Figure 8:**
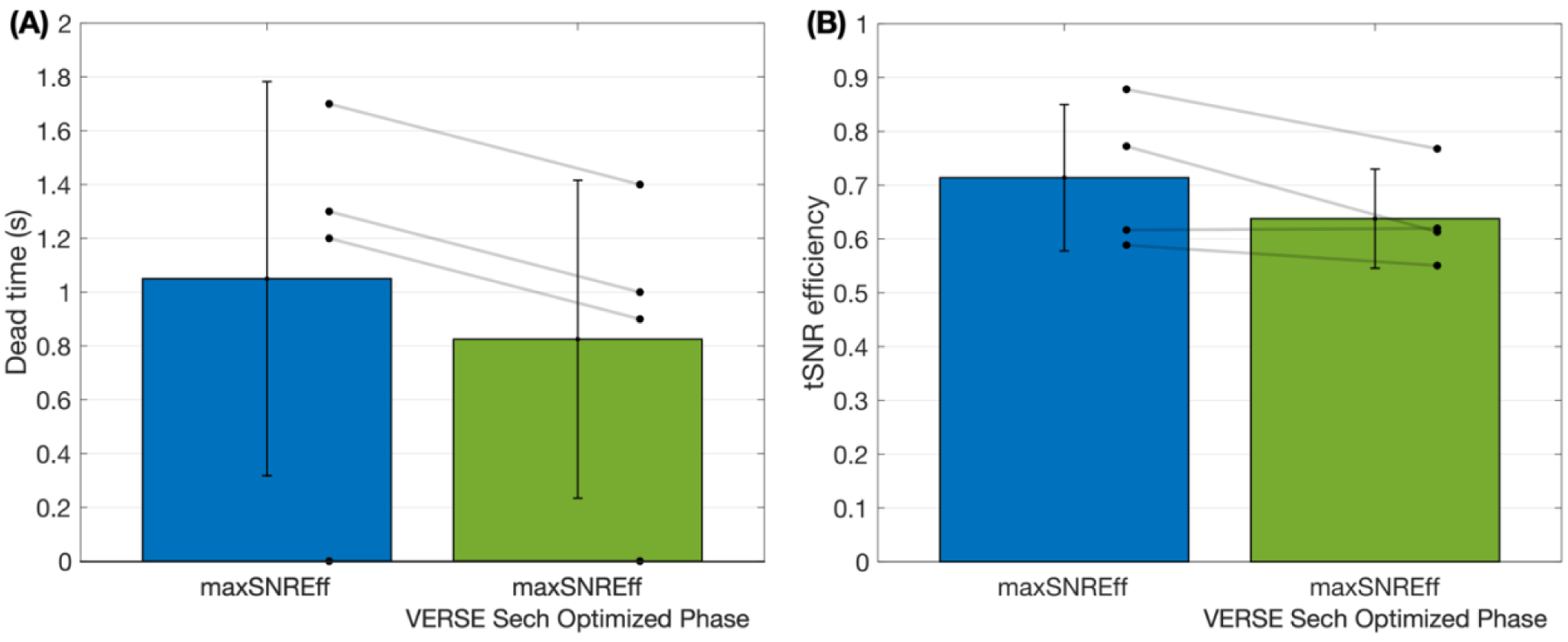
The results of the in vivo SNR efficiency comparison between the max SNR efficiency (maxSNREff) protocol with the standard sech pulse (blue) and the VERSE sech pulse with optimized phase waveform (green). (A) The minimum added sequence deadtime achieved for each protocol, subject to the 1st level SAR limits. (B) The subject-level mean gray matter SNR efficiencies for each protocol, calculated as the temporal SNR divided by the square-root of the TR. Both protocols achieved the target label duration of 1800 ms. The bar graphs show the mean and standard deviation across subjects. Individual subject values are plotted as dots. The TRs and tSNR efficiencies were not significantly different between these two protocols.

### 3.4. High-resolution demonstration

The high-resolution data (1.95x1.95x4 mm^3^) is shown in Figure 9 alongside the standard resolution data (3.44x3.44x5 mm^3^). Both scans used the max SNR efficiency PCASL parameters with the standard sech BGS pulse. The scan time in both cases was 4 minutes and, despite a reduction in voxel volume by a factor of 3.9, the high-resolution data still demonstrates high SNR but with greatly improved spatial information.

**Figure 9:**
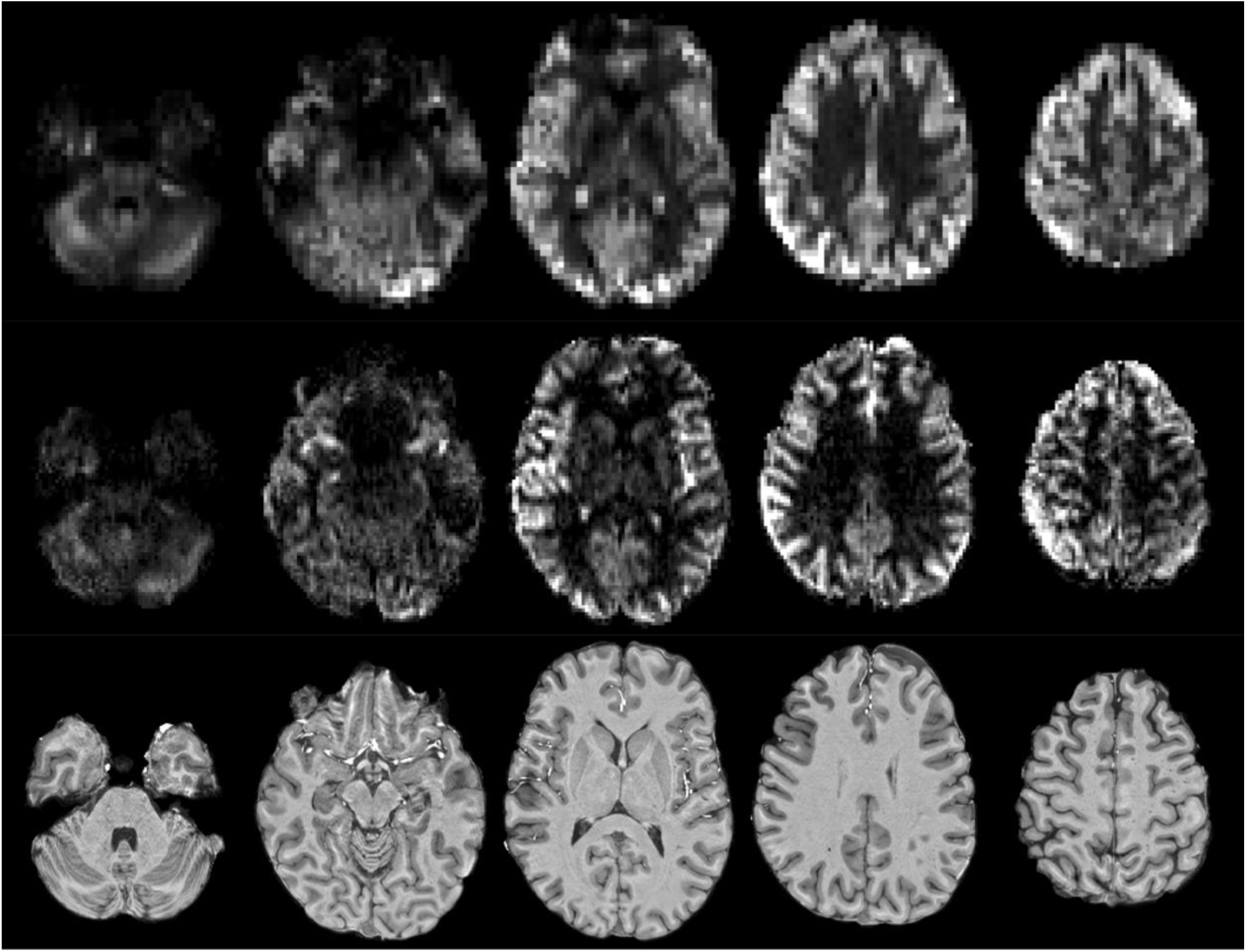
Qualitative comparison of an example low (top row, 3.44 x 3.44 x 5 mm^3^) and high (middle row, 1.95 x 1.95 x 4 mm^3^) spatial resolution perfusion weighted images at several slices in a single subject. The slices are approximately matched in location. The bottom row shows matching slices of the corresponding MPRAGE structural image.

## 4. Discussion

The primary novel contribution of this study was to optimize the PCASL preparation to maximize SNR efficiency in SAR-constrained scenarios, such as those found at 7 T. A previous study^12^ optimized the PCASL RF and gradient parameters to maximize labeling efficiency over a large range of B_1_^+^ and B_0_, thus making the PCASL labeling less sensitive to these variations. However, this inevitably leads to PCASL settings with high RF power (short TR_PCASL_ and high nominal B_1_^+^_ave_), requiring long TRs due to SAR limits. Here, we instead explicitly correct for variations in B_1_^+^ and B_0_ in the feeding vessels at the labeling plane with fast calibration scans (total scan time 24 s). This allowed us to use the available degrees-of-freedom for choosing the PCASL parameters to instead maximize SNR efficiency, where labeling efficiency is balanced against the total scan RF power, yielding a 126% improvement in SNR efficiency in vivo compared to 7T PCASL settings from the literature.^22^

The optimized max SNR efficiency parameters maintained a high labeling efficiency whilst achieving a greatly reduced RF power (2.4-4.9 times lower than the comparison protocols) by combining a low B_1_^+^_ave_ with a low G_ave_. Although reducing B_1_^+^_ave_ reduces the labeling efficiency of fast flowing spins, lowering G_ave_ counteracts this effect.^17,42^ Nevertheless, lowering B_1_^+^ave reduces the velocity range that experiences high labeling efficiency,^17^ as seen in Figure 3. Despite this, the max SNR efficiency parameters achieved only modestly lower labeling efficiency than the other protocols for the pulsatile flow waveform and off-resonance range considered here (0.72 vs 0.79-0.82). This pulsatile flow waveform, which has a mean velocity of about 40 cm/s, is representative of typical peak flow in the internal carotid arteries of young healthy volunteers,^43^ with the represented velocities higher than those of vertebral arteries and healthy older subjects. Since the SNR efficiency of the max SNR efficiency parameters is even higher at velocities lower than 40 cm/s compared to the literature parameters, this should increase the SNR efficiency advantage of this protocol even more in these populations. However, the potentially large benefits of the max SNR efficiency settings still need to be validated in clinical populations in future studies. Additionally, where the blood flow velocities differ substantially from those used here, the PCASL parameters would, ideally, be reoptimized to maximize performance.

The transmit coil 1 s RF power limit greatly restricted the label durations of the higher SAR comparison protocols, ultimately leading to a different comparison than originally intended: we had aimed to use a 1800 ms label duration for each protocol, extending the TR with more deadtime for the higher RF power protocols to stay within 1st-level SAR limits. However, although not experimentally feasible with our MRI system, our simulation results for equal label durations of 1800 ms suggest that the max SNR efficiency protocol would have 55%, 34%, and 22% higher SNR efficiency in vivo than the max labeling efficiency, literature, and literature with VERSE protocols, respectively. Although smaller than the results achieved in vivo, they still represent a large improvement in SNR efficiency.

The static tissue response of the max SNR efficiency parameters had a central lobe that extended 1.78 cm beyond the center of the labeling plane, 8.6 mm less than that of the literature protocol. This means the labeling plane can be placed closer to the imaging volume without causing artifacts in the perfusion weighted images. This is especially beneficial for whole brain imaging at UHF, because B_1_^+^ typically drops off quickly towards the bottom of the brain with standard head-only transmit coils, making it unfeasible to use labeling plane positions lower than that used here.

Previous 7 T PCASL work^22^ explored optimizing the adiabatic BGS pulses to achieve improved inversion efficiency. As might be expected, greater inversion efficiency across a broad range of B_1_^+^ and B_0_ was achieved with greater RF power, though with diminishing returns. However, because the effect of RF power on the TR and SNR efficiency was not incorporated into the optimization, the highest RF power pulse was used. Here, we instead explored whether a lower RF power inversion pulse could further increase SNR efficiency, even though its inversion efficiency was slightly lower than the original pulse. Unfortunately, even though this lower RF power pulse enabled shorter TRs, the in vivo SNR efficiency decreased by 11%, in contrast to simulations. This could be for several reasons, including: a lower blood inversion efficiency than expected, leading to more ASL signal loss; or it could be because the decreased inversion efficiency led to poorer BGS and more physiological noise. We also explored HSn^44^ pulses for n≤8 using simulations, but found these had lower inversion efficiency than the optimized-phase VERSE sech pulse, even when their RF power was matched to the standard sech pulse.

Another set of PCASL parameters from the max SNR efficiency optimization (RF flip angle=7.5°, B_1_^+^_ave_=0.7 µT, T_RF_=350 µs, TR_PCASL_=700 µs, G_max_=8 mT/m, G_ave_=0.3 mT/m) were found to have a theoretical SNR efficiency only 0.33% lower than those in Table 1. However, because these settings use a shorter TR_PCASL_ (700 µs versus 1060 µs), they will be more robust to cases where the off-resonance in the feeding arteries at the labeling plane deviates by more than ±50 Hz (Figure S7). Nevertheless, these settings would have exceeded the transmit coil 1 s power limit by a small amount for all but one subject in this study.

Although we used T_1_^+^ and T_2_ values appropriate for blood at 7 T for the static tissue Bloch simulations, the static tissue response for the protocols in Table 1 do not change greatly when using values more appropriate for brain tissue (T1=1500 ms and T2=50 ms,^22^ results not shown), therefore, we do not expect that this choice markedly affected the results of the optimization.

Another approach to reduce SAR, that was not explored here, is to increase the RF duty cycle beyond 50%.^19^ However, this can increase the level of RF amplifier drift, as recently reported.^45^ This effect modulates B_1_^+^_ave_ during the labeling train and can also result in SAR limits being reached unexpectedly, with the scan being prematurely stopped by the real-time SAR monitoring system. We investigated the level of RF amplifier drift using the max SNR efficiency protocol with different RF duty cycles to evaluate this effect ourselves. We observed a drift of 7.5% across a 1.8 s label duration at 50% duty cycle, but this increased to 10.7% at the maximum possible duty cycle of 68% (Figure S8). In future studies, the correction proposed by Aghaeifar et al.^45^ could be used to enable the use of higher RF duty cycles, further reducing SAR.

At 3T, long label durations are expected to be more SNR efficient than the standard 1.8 s,^46^ and recent results suggest this is still the case at 7 T, where the minimum TR is SAR constrained.^25^ Therefore, this could be another approach to further increase the SNR efficiency of 7 T PCASL perfusion imaging. Indeed, when we simulated the SNR efficiency of the max SNR efficiency protocol for a range of label durations from 100 - 8000 ms, sampled every 100 ms, by multiplying the righthand side of **Error! Reference source not found.** by the expected perfusion signal (2015 consensus paper model^3^, PLD=1800 ms, perfusion=50 mL/100 g/min, T_1_^+^=2.1 s), the theoretical optimal LD duration is 4100 ms when assuming the mean in vivo B_1_^+^.

## 5. Conclusions

We optimized the PCASL parameters to maximize SNR efficiency for whole brain perfusion imaging at 7 T by balancing labeling efficiency with the total RF power. By using a low B_1_^+^_ave_, G_ave_, and G_max_, with a long TR_PCASL_, we achieved a significant reduction in RF power while maintaining high labeling efficiency. This resulted in a 126% improvement in in vivo SNR efficiency compared to existing 7T PCASL settings and allowed for longer label durations and minimized deadtime.

## 6 Acknowledgements

We thank Dr. Thijs de Buck for sharing his experience with 7 Tesla scanning, Dr. James Kent for his advice on B_1_^+^ mapping, Drs. Philipp Ehses and Tony Stöcker for sharing their 3DREAM sequence, Dr. Li Zhao for sharing the carotid artery velocity waveform used in this work, and Dr. Yulin Chang for assistance with implementing dynamic linear B_0_-shimming. The authors also thank Dr. Brian Hargreaves for making his minimum time VERSE code freely available (http://mrsrl.stanford.edu/~brian/mintverse/), as we built on this to develop our minimum SAR VERSE code.

## 7. Data availability statement

The code for simulating and optimizing the PCASL pulse train parameters and the background suppression adiabatic inversion pulses is available at (**Zenodo link**). The VERSE algorithm we developed for this work is available at (**Zenodo link**). Unfortunately, we are currently unable to share the full in vivo data due to data protection issues, although the Wellcome Centre for Integrative Neuroimaging is actively working on a solution to this. Nevertheless, the summary data and analysis code that underly the figures in this manuscript are available at (**Zenodo link**).

## 8. Funding information

J.G.W. and T.W.O. were supported by a Sir Henry Dale Fellowship jointly funded by the Wellcome Trust and the Royal Society (220204/Z/20/Z). Y.J. is supported by National Natural Science Foundation of China (62401535). We thank the UK BBSRC (grant number BB/W019582/1) and the NIHR Oxford Health Biomedical Research Centre (NIHR203316) for support. The views expressed are those of the authors and not necessarily those of the NIHR or the Department of Health and Social Care. J.G.W. acknowledges Linacre College (Oxford) for their support. The Wellcome Centre for Integrative Neuroimaging is supported by core funding from the Wellcome Trust (203139/Z/16/Z and 203139/A/16/Z). For the purpose of open access, the author has applied a CC BY public copyright license to any author-accepted manuscript version arising from this submission.

## Footnotes

* WM perfusion is 3-4 times lower than GM perfusion and has longer arterial transit times (ATTs)7,8, resulting in much lower SNR.

## Notes

### Competing Interest Statement

The authors have declared no competing interest.

