## Supporting Information for "SNR-Efficient Whole-Brain Pseudo-Continuous Arterial Spin Labeling Perfusion Imaging at 7 Tesla"

### 1. Supporting Information 1 - Reduced SAR background suppression

We applied our minimum SAR VERSE algorithm to a sech pulse (duration = 10.24 ms, β = 763.99 rad/s, $B_1^+{}_{max}$ = 20 μT), optimizing μ for the resulting reformatted RF pulse in the same way as described above. The maximum VERSE amplitude compression/dilation factor was limited to 1.43, chosen to give a SAR reduction of 25%. This SAR reduction is similar to that of the PCASL pulses and limits the reduced off-resonance performance seen at higher SAR reduction levels.[34]

The phase waveform of the VERSE sech pulse was then further optimized to maximize the inversion efficiency over the same $B_1^+$ and $B_0$ ranges as before (±50% and ±500 Hz, respectively) using MATLAB's sequential quadratic programming algorithm (fmincon). The individual phase samples in the $B_1^+$ waveform (sampled every 10 μs) were treated as free parameters to optimize within the range ±pi rad.

A multi-TI saturation-inversion-recovery experiment (single sagittal slice with EPI readout) was performed in a single subject to validate the performance of the standard sech pulse and VERSE phased-optimized sech pulse. The time between the saturation and inversion was 8 s, and the TIs were [100, 150, 220, 320, 480, 710, 1050, 1550, 2290, 3380, 5000] ms, i.e. an exponential sampling pattern. After motion and distortion correction, the multi-TI data was simultaneously fit for $T_1$ and inversion efficiency for each dataset.

In vivo perfusion data using the max SNR efficiency PCASL parameters and the resulting phase-optimized VERSE sech pulse were also acquired in the same four volunteers using an otherwise matched standard resolution protocol to that described in section 2.3.2 of the paper.

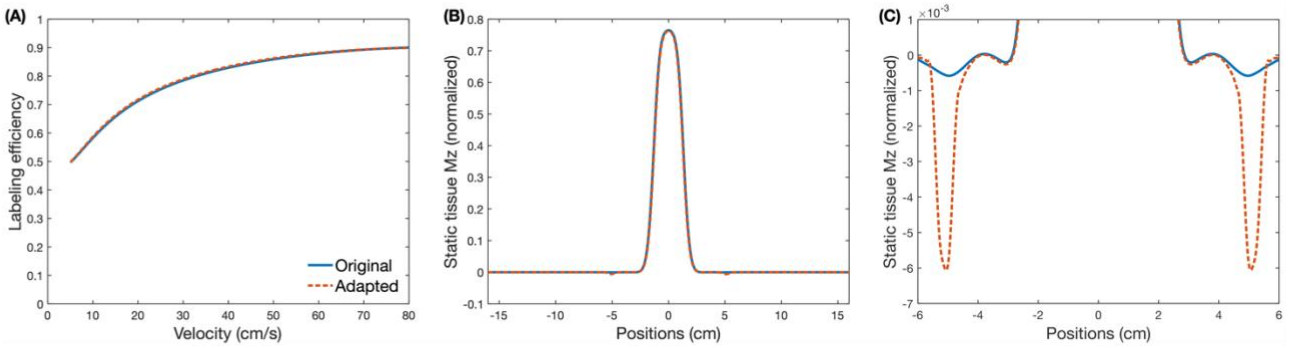


Supporting Information Figure S1: Comparing the labeling efficiency (A) and static tissue response (B,C) of Zhao et al.'s PCASL parameters ("Original") and the slightly adapted parameters used in this work ("Adapted"). (A) shows the labeling efficiency for constant velocity plug flow across a range of velocities for on-resonant spins. This graph demonstrates that the labeling efficiency of the two sets of parameters is almost identical. (B) shows the static tissue difference Mz after a labeling duration of 1800 ms and convolution with the slice profile of the readout excitation pulse. The static tissue response is almost identical between the two sets of parameters, though the zoomed in view of (B) shown in (C) highlights that there is a small aliased labeling plane perturbation with the adapted set of parameters (note the small y-axis range in (C)). Use of Zhao et al.'s aliased labeling plane suppression formula with the increased $TR_{PCASL}$ would have required an increase of $G_{max}$ to 6.08 mT/m, which would violate gradient slew rate constraints on our system. Therefore, in the interest of keeping the settings as close as possible to the original values, and because the static tissue perturbation was considered unlikely to grossly influence the in vivo comparison, no further changes were made.

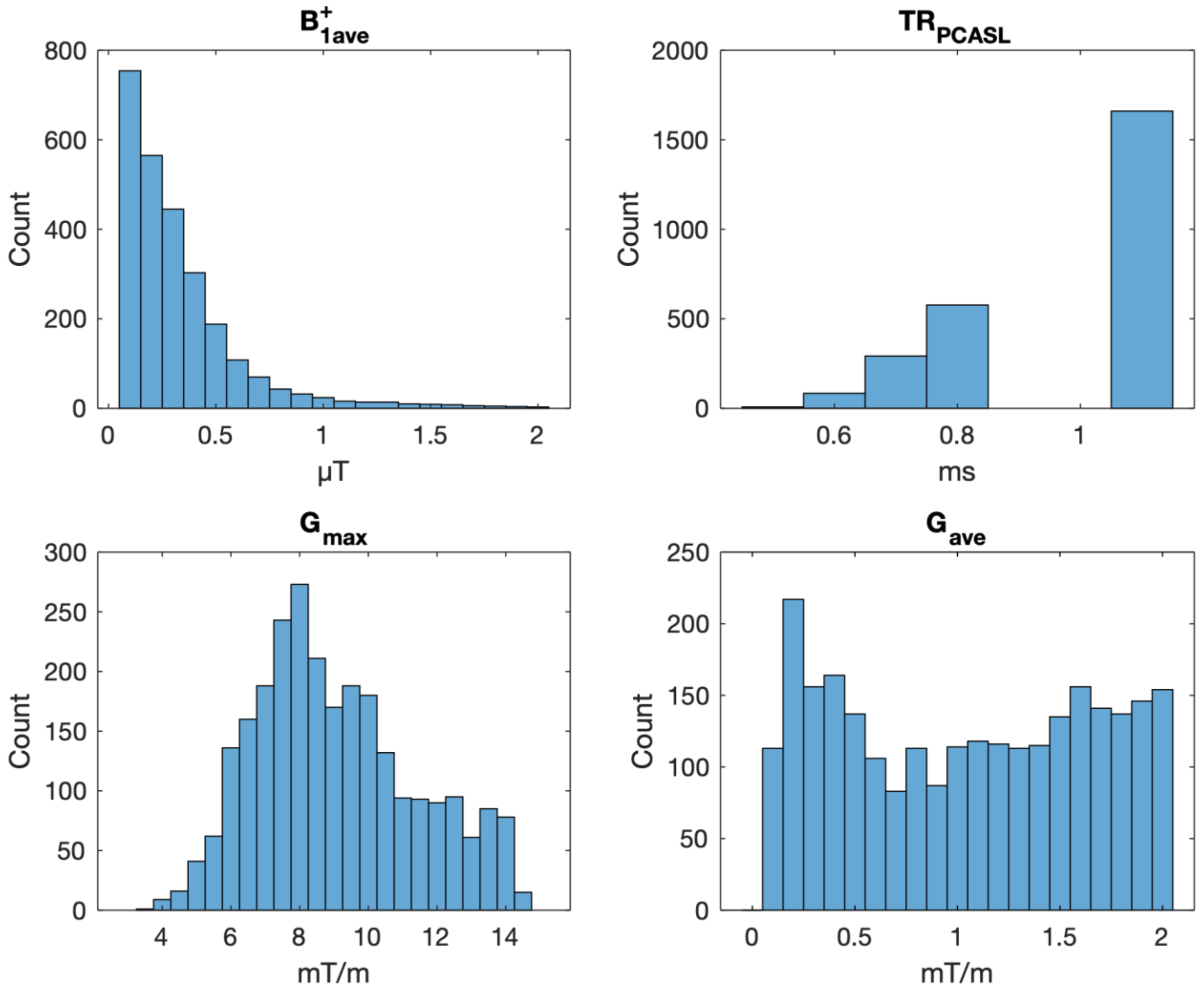


Supporting Information Figure S2: Histograms of individual PCASL parameters for the parameter combinations that did not exceed gradient slew rate limits and did not have static tissue perturbation of more than 0.1% of $|M_0|$ beyond 1.8 cm from the labeling plane. In general, there was an increasing probability of lower $B_1^+{}_{ave}$ and longer $TR_{PCASL}$, with the distribution for $G_{max}$ peaking at 8 mT/m. These trends are because lower $B_1^+{}_{ave}$ values have a decreased effect on static tissue, while a higher $G_{max}$ will also have a smaller slice width and so less effect on static tissue beyond 1.8 cm. However, as $G_{max}$ increases, slew rate issues become more likely, hence the decreased frequency beyond 8 mT/m. Additionally, a longer $TR_{PCASL}$ means that gradient slew rate issues are less likely. There was a less clear pattern with the surviving $G_{ave}$ values due to the complex interactions of this parameter with others in determining the static tissue perturbations.

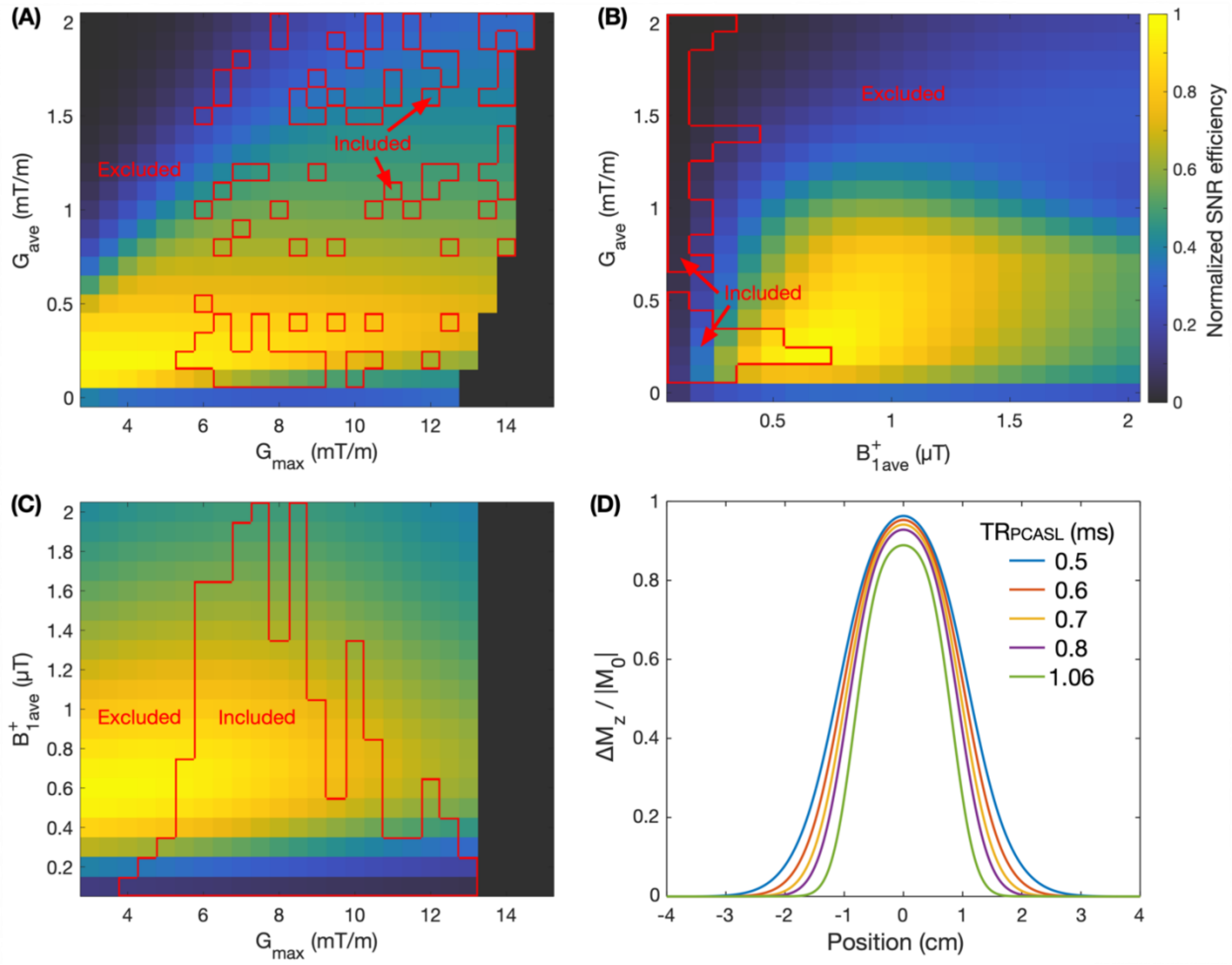


Supporting Information Figure S3: In (A-C), the simulated SNR efficiencies, normalized to the highest SNR efficiency, are shown across the full range of two PCASL parameters with the other parameters fixed at the max SNR efficiency values (Table 1). The zero values in (A) and (C) are parameter combinations that exceeded gradient slew rate limits and were not simulated. The red outlines in (A-C) highlight the parameter combinations that satisfied the static tissue perturbation constraint and were thus included in the optimization. (D) shows the central portion of the simulated ΔMz static tissue response to a 1800 ms long PCASL train for the optimized maximum SNR efficiency parameters for the considered $TR_{PCASL}$ values, demonstrating a decreasing central lobe width as $TR_{PCASL}$ increases.

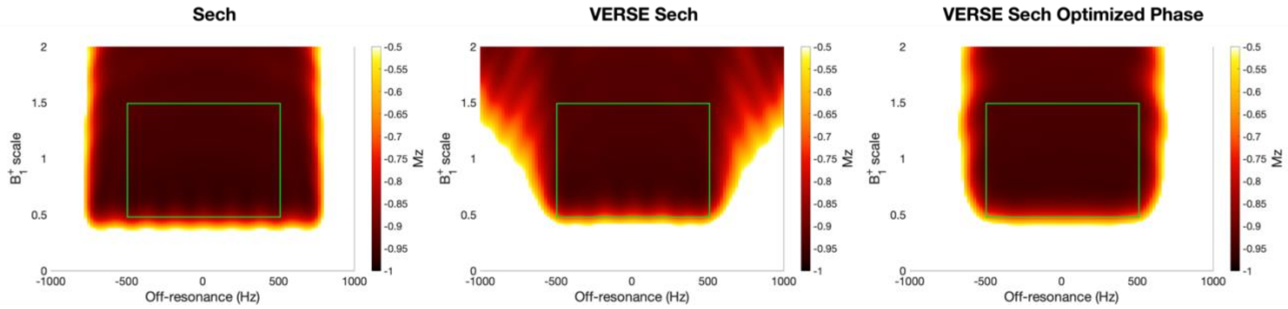


Supporting Information Figure S4: The same inversion maps as in Figure 7, but windowed differently to highlight the differences in performance within the target $B_1^+$ and $B_0$ range ($\Delta B_1^+ = \pm 50\%$, $\Delta B_0 = \pm 500$ Hz, green box).

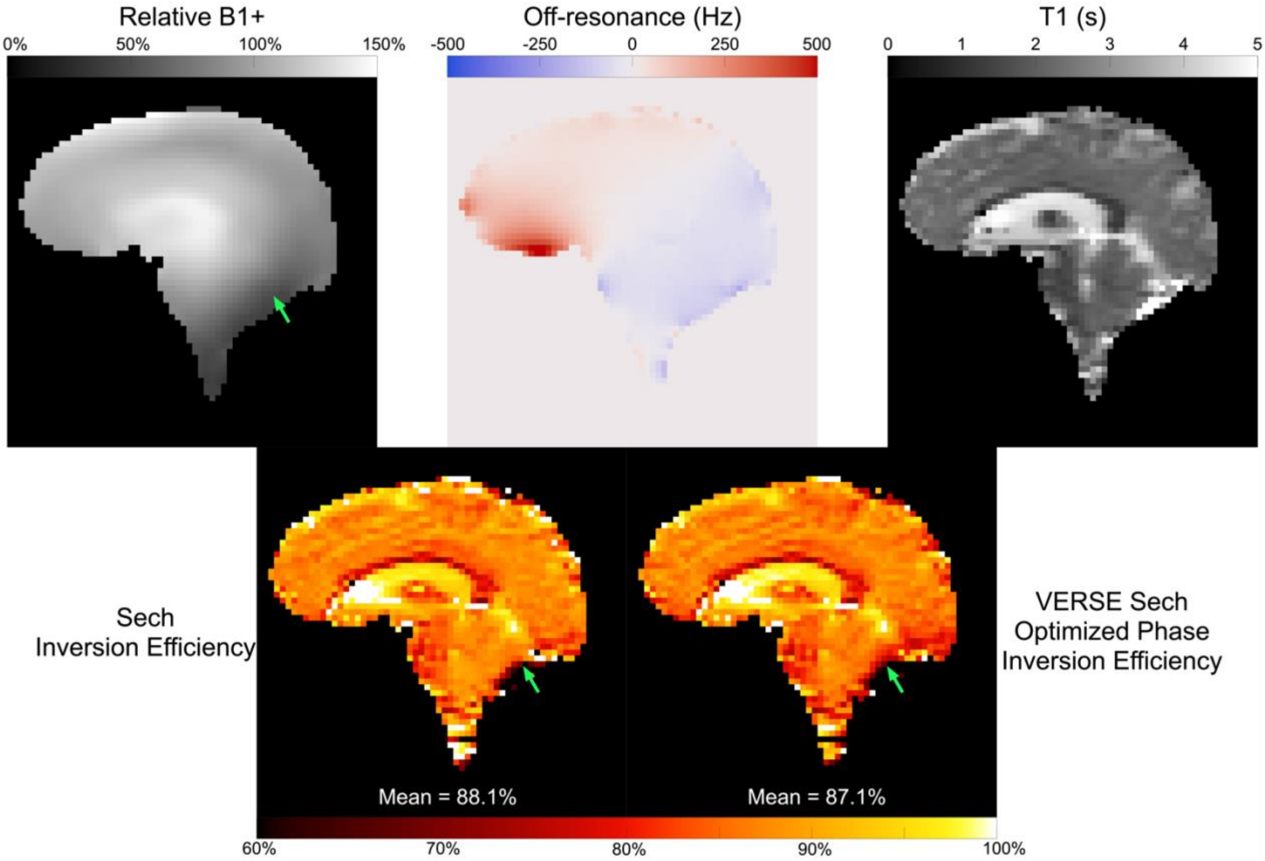


Supporting Information Figure S5: The results of the single-slice, multi-TI, saturation-inversion-recovery experiment used to measure the inversion efficiency of each inversion pulse in vivo. The relative $B_1^+$ were calculated using the 3DREAM sequence and is relative to the reference voltage. The off-resonance map was measured using a single-slice dual echo GRE sequence. The $T_1$ and inversion efficiencies were then jointly estimated from the multi-TI saturation-inversion-recovery datasets. The green arrow highlights an area of low $B_1^+$ that leads to lower inversion efficiency with the VERSE sech optimized phase pulse than with the standard sech pulse.

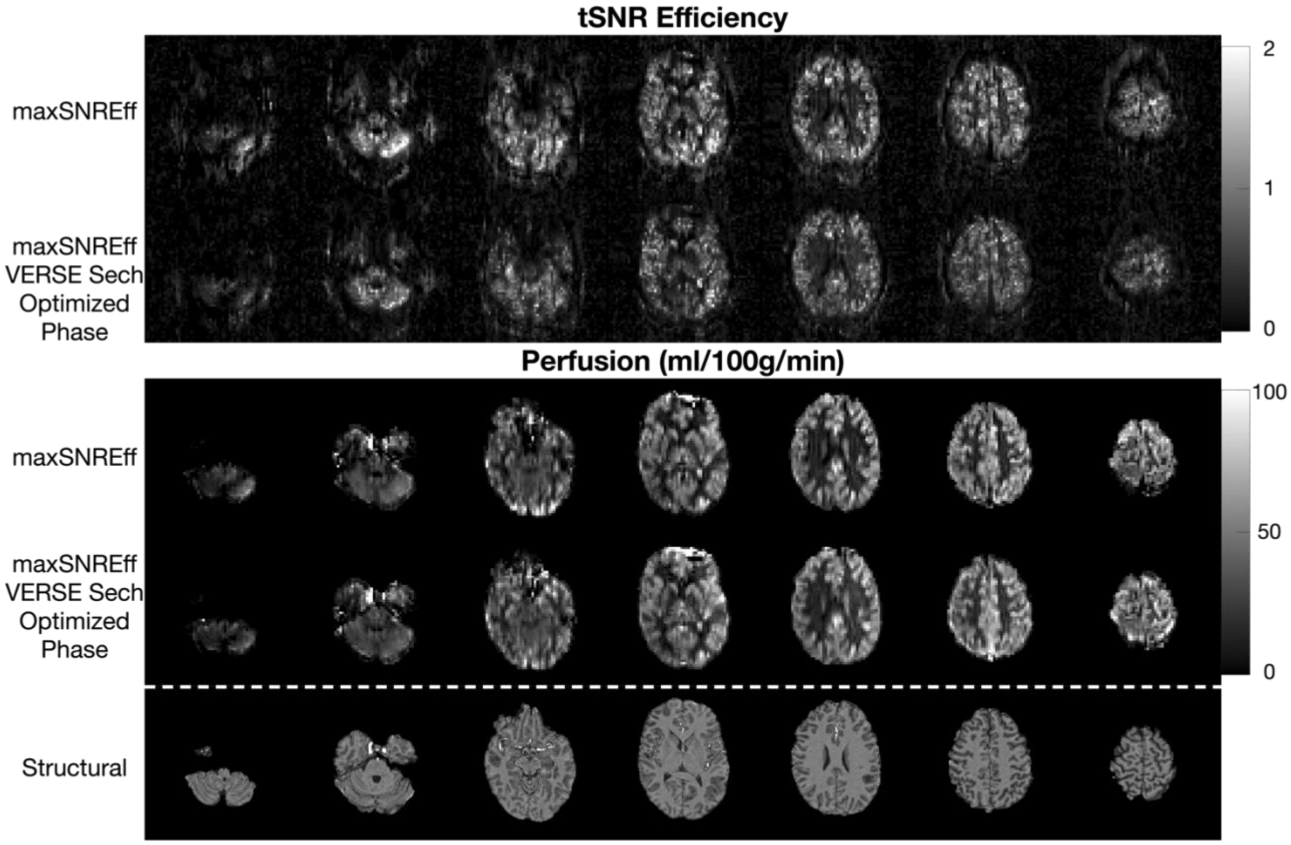


Supporting Information Figure S6: The tSNR efficiency and CBF maps in one subject from the comparison between the max SNR efficiency PCASL parameters with either the sech or VERSE sech with optimized phase background suppression pulses. Each row shows seven slices from each protocol, with the corresponding slices from the MPRAGE structural scan shown in the bottom row.

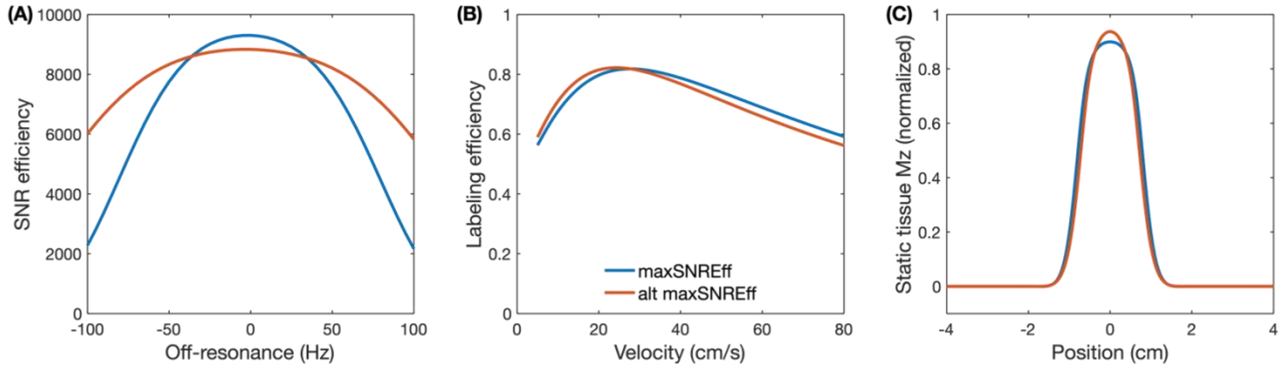

Supporting Information Figure S7: Comparing the SNR efficiency across off-resonance (A), plug-flow on-resonant labeling efficiency across velocities (B), and the static tissue response (C) of the max SNR efficiency PCASL parameters (maxSNREff) and the alternative PCASL parameters (alt maxSNREff) that were found to have a theoretical SNR efficiency only 0.33% lower than the max SNR efficiency parameters. The max SNR efficiency parameters have higher mean SNR efficiency for the -50 Hz - 50 Hz range, but the alternative parameters are more robust to larger off-resonance ranges due to the shorter $TR_{PCASL}$ used, so might be useful in other studies where dynamic $B_0$ shimming is not available or does not sufficiently correct for off-resonance. The on-resonant labeling efficiency and static tissue response are similar.

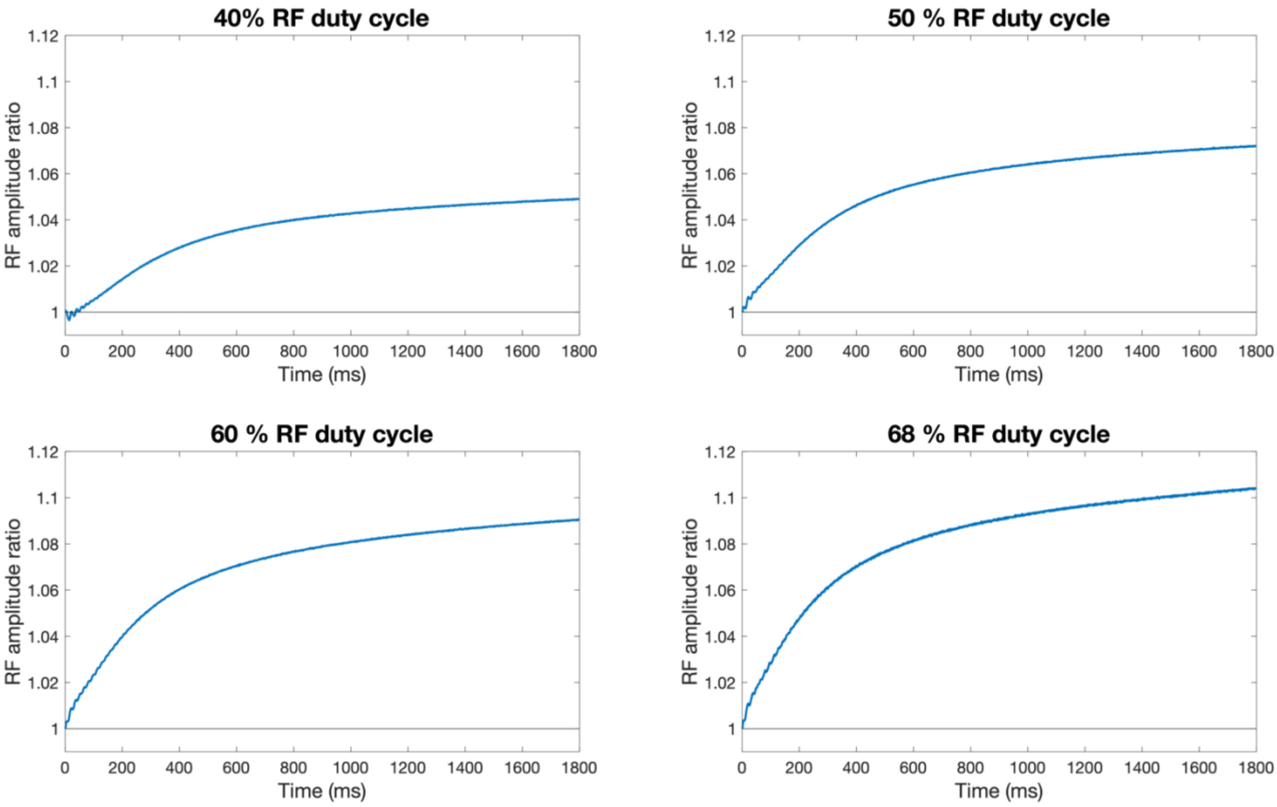


Supporting Information Figure S8: The measured PCASL RF amplifier drift, averaged over all TRs of a 4-minute scan, for the max SNR efficiency protocol using different RF duty cycles. The maximum drift across the 1800 ms label duration in each case was 5.2%, 7.5%, 9.3%, and 10.7% for duty cycles of 40%, 50%, 60%, and 68%, respectively.
